# Microglia Proliferation Underlies Synaptic Dysfunction in the Prefrontal Cortex: Implications for the Pathogenesis of HIV-1-Associated Neurocognitive and Affective Alterations

**DOI:** 10.1101/2023.01.20.524942

**Authors:** Hailong Li, Kristen A. McLaurin, Charles F. Mactutus, Rosemarie M. Booze

## Abstract

Microglia, which are productively infected by HIV-1, are critical for brain development and maturation, as well as synaptic plasticity. The pathophysiology of HIV-infected microglia and their role in the pathogenesis of HIV-1-associated neurocognitive and affective alterations, however, remains understudied. Three complementary aims were undertaken to critically address this knowledge gap. First, the predominant cell type expressing HIV-1 mRNA in the dorsolateral prefrontal cortex of postmortem HIV-1 seropositive individuals with HAND was investigated. Utilization of a combined RNAscope multiplex fluorescent and immunostaining assay revealed prominent HIV-1 mRNA in microglia of postmortem HIV-1 seropositive individuals with HAND. Second, measures of microglia proliferation and neuronal damage were evaluated in chimeric HIV (EcoHIV) rats. Eight weeks after EcoHIV innoculation, enhanced microglial proliferation was observed in the medial prefrontal cortex (mPFC) of EcoHIV rats, evidenced by an increased number of cells co-localized with both Iba1+ and Ki67+ relative to control animals. Neuronal damage in EcoHIV infected rats was evidenced by pronounced decreases in both synaptophysin and post synaptic density protein 95 (PSD-95), markers of pre-synaptic and post-synaptic damage, respectively. Third, regression analyses were conducted to evaluate whether microglia proliferation mechanistically underlies neuronal damage in EcoHIV and control animals. Indeed, microglia proliferation accounts for 42-68.6% of the variance in synaptic dysfunction. Collectively, microglia proliferation induced by chronic HIV-1 viral protein exposure may underlie the profound synaptodendritic alterations in HIV-1. Understanding how microglia are involved in the pathogenesis of HAND and HIV-1-associated affective disorders affords a key target for the development of novel therapeutics.

## INTRODUCTION

Individuals living with human immunodeficiency virus type 1 (HIV-1) are treated with combination antiretroviral therapy (cART), which successfully establishes virological control in the periphery (Autran et al., 1997). The central nervous system (CNS), however, is infiltrated with infected monocytes and CD4+ cells within days of HIV infection (Koenig et al., 1986; Whitney et al., 2014). Consequently, nearly half of HIV-1 seropositive individuals are afflicted with debilitating neurocognitive impairments (NCI; Wang et al., 2020; Zenebe et al., 2022) and/or affective (Campos et al., 2010; Kamat et al., 2013) alterations. Indeed, HIV-1-associated neurocognitive disorders (HAND) are characterized by deficits in attention, memory, and executive function (Heaton et al., 2011), whereas affective alterations include both apathy and depression (for review, Cysique & Brew, 2019). The broader implications of HIV-1-associated neurocognitive and affective alterations cannot be understated, as they are associated with deficits in everyday functioning (e.g., Vance et al., 2011; Kamat et al., 2012) and decreased medication adherence (Becker et al., 2011; Panos et al., 2014). Given the prevalence and implications of HIV-1-associated neurocognitive and affective alterations, there remains a fundamental need to better characterize the pathogenesis of these disorders.

Infected peripheral cells (i.e., monocytes and CD4+ cells), carrying either the integrated HIV-1 provirus or unintegrated circular HIV-1 DNA, migrate across the blood-brain barrier (Veenstra et al., 2017) infecting predominantly microglia (e.g., Ko et al., 2019; Li et al., 2021a). Microglia, the brain-resident macrophages, account for 5-20% of the total cells in the adult brain (Polazzi et al., 2010), whereby they are characterized by a small cell body and very thin ramified processes. Under homeostatic conditions, microglia are fundamentally involved in the development and maintenance of neural circuits (e.g., Paolicelli et al., 2011; Squarzoni et al., 2014) and synaptic plasticity (e.g., Ji et al., 2013; Raghuraman et al., 2019). Infection with HIV-1, however, likely alters the morphology and physiology of microglia. Indeed, microglial activation in chronically infected HIV-1 seropositive individuals has been evidenced by increased translocator protein binding (Garvey et al., 2014); activation which is inversely associated with neurocognitive function (Rubin et al., 2018). Furthermore, microglia serve as a viral reservoir for HIV-1 (for review, Wallet et al., 2019; Ko et al., 2019; Li et al., 2021a). Additional investigations, however, are necessary to better understand the pathophysiology of HIV-infected microglia and their role in the pathogenesis of HIV-1-associated neurocognitive and affective alterations.

Although the etiology of HIV-1-associated neurocognitive and affective alterations is multifaceted, compelling evidence from HIV-1 seropositive individuals supports the fundamental role of neuronal injury (Moore et al., 2006) and synaptic dysfunction (Gelman & Nguyen, 2010; Desplats et al., 2013; Weiss et al., 2021). The functional implications of synaptodendritic alterations has also been demonstrated, as neuronal and synaptic damage is significantly correlated with individual neurocognitive domains and global neurocognitive impairment in HIV-1 seropositive individuals (Moore et al., 2006; Levine et al., 2016). Critical to the current hypothesis is the finding that microglial activation, indexed using cerebral spinal fluid (CSF) concentrations of soluble TREM2, is associated (*r*=0.62) with a measure of neuronal injury (i.e., CSF Neurofilament Light Protein; Gisslén et al., 2018) in HIV-1 seropositive individuals. The ethical and technical constraints of studying CNS disorders, and their pathophysiology, in humans necessitates valid preclinical biological systems.

Indeed, preclinical biological systems have been instrumental in recapitulating and expanding our understanding of the pathophysiology of HIV-1-associated neurocognitive and affective alterations. Consistent with observations in HIV-1 seropositive individuals, preclinical biological systems, including Tat transgenic (Tg) mice (e.g., Fitting et al., 2013), gp120 Tg mice (e.g., Speidell et al., 2020) and the HIV-1 Tg rat (e.g., Festa et al., 2020), exhibit pronounced decreases in synaptic density relative to their control counterparts. Three-dimensional reconstruction analyses (Li et al., 2020) have more intricately described HIV-induced alterations in neuronal and dendritic spine morphology (e.g., Roscoe et al., 2014; McLaurin et al., 2019; Li et al., 2021a; McLaurin et al., 2022). More specifically, continuous measures of dendritic spine morphology (e.g., backbone length, head diameter, neck diameter) support a population shift towards a more immature dendritic spine phenotype in both the HIV-1 Tg (Roscoe et al., 2014; McLaurin et al., 2019) and chimeric HIV (EcoHIV; Li et al., 2021; McLaurin et al., 2022) rats. Preclinical biological systems, therefore, afford an opportunity to evaluate the generalizability of the association between microglial activation and synaptic dysfunction.

In light of previous observations, three aims were untaken to critically address this fundamental knowledge gap. First, using a combined RNAscope multiplex fluorescent and immunostaining assay, the predominant cell type expressing HIV-1 mRNA in the dorsolateral prefrontal cortex (dlPFC) of postmortem HIV-1 seropositive individuals with HAND was investigated. Second, measures of microglia proliferation and neuronal damage were evaluated in chimeric HIV (EcoHIV) rats eight weeks after inoculation. Third, regression analyses were conducted to evaluate whether microglia proliferation mechanistically underlies neuronal damage in EcoHIV and control animals. Taken together, the present study affords an opportunity to expand upon our understanding of the pathophysiology of HIV-infected microglia and their role in the pathogenesis of HIV-1-associated neurocognitive and affective alterations.

## MATERIALS AND METHODS

### Experiment #1: HIV-1 Seropositive Individuals with HIV-1-Associated Neurocognitive Disorders

#### Autopsy Human Brain Tissues

The National NeuroAIDS Tissue Consortium (NNTC) provided paraformaldehyde-fixed autopsy human brain tissue from the dorsolateral prefrontal cortex (dlPFC; Brodmann’s Area 9 (Brodmann, 1909)) of nine HIV-1 seropositive individuals with HAND. Brain tissue was from individuals between 55 and 74 years of age diagnosed with symptomatic HAND at any point during study participation. The NNTC Data Coordinating Center approved the specimen application (Request # R703).

#### Combined RNAscope Multiplex Fluorescent and Immunostaining Assay

A combined RNAscope multiplex fluorescent and immunostaining assay was utilized to evaluate the predominant cell type expressing HIV-1 mRNA in post-mortem HIV-1 seropositive individuals with HAND. RNAscope *in situ* hybridization procedures were described in detail by (Li et al., 2018), albeit with minor modifications for the present study. In brief, a cryostat was used to cut 50 µm thick sections of 4% paraformaldehyde-fixed human brain tissue. Subsequently, brain tissues were transferred and mounted onto slides, which were boiled in 1×Target retrieval buffer for 5 min. Sections were hybridized at 40°C with a probe for HIV-1 mRNA (Advanced Cell Diagnostics Inc., Newark, CA, USA; Cat. No. 444061) for 2 hours. Next, sections were incubated at 40°C for 30 min with Opal 570 Dyes (Akoya Biosciences, Marlborough, MA, USA; Cat. No. FP1488001KT). Sections were subsequently incubated overnight with rabbit monoclonal anti-Iba1 antibody (Abcam, Cambridge, United Kingdom; Cat. No. 178847) at 4°C. Opal 520 dyes (Akoya Biosciences, Cat. No. FP1487001KT) were applied to sections for 10 min at room temperature. Slides were mounted with Pro-Long Gold Antifade reagent (Invitrogen, Carlsbad, CA, USA) and imaged at 60× using a Nikon TE-2000E confocal microscope system.

#### Immunofluorescence Staining

50 µm thickness coronal slices were incubated overnight at 4°C with Alexa Fluo 488 rabbit monoclonal anti-synaptophysin antibody. Fluorescent images were captured using a Nikon D-Eclipse C1 Confocal microscope. Fluorescence signal intensity was analyzed using NIS-Elements BR3.10 software.

### Experiment #2: EcoHIV Infection of Rats

#### Animals

Adult Fischer F344/N rats (*n*=12; Male, *n*=8; Female, *n*=4) were procured from Envigo Laboratories Inc. (Indianapolis, IN, USA) and randomly assigned to receive stereotaxic injections of either the EcoHIV lentivirus (*n*=6; Male, *n*=4; Female, *n*=2) or saline (i.e., control; n=6; Male, *n*=4; Female, *n*=2) into the cortex (0.8 mm lateral and 1.2 mm anterior to Bregma, 2.5 mm deep). Construction of the EcoHIV virus and methods utilized for stereotaxic surgeries were previously described (Li et al., 2021a).

Rats were pair-housed in AAALAC-accredited facilities at the University of South Carolina according to guidelines established by the National Institutes of Health. Environmental conditions of the animal facility were targeted at 12h:12h light/dark cycle with lights on at 700 h (EST), 21°± 2°C, and 50% ± 10% relative humidity. *Ad libitum* access to rodent chow (20/20X, Harlan Teklad, Madison, WI, USA) and water were provided throughout the duration of the study. All procedures were approved by the Institutional Animal Care and Use Committee (IACUC) of the University of South Carolina (Federal Assurance #D16-00028).

#### Immunofluorescence Staining

Following induction with the anesthetic sevofluorane (Abbot Laboratories, North Chicago, IL, USA), EcoHIV and control animals were transcardially perfused with 4% paraformaldehyde. A Vibratome (PELCO easiSlicer(tm) Vibratory Tissue Slicer, Ted Pella, CA, USA) was used to cut coronal sections of brain tissue with a thickness of 100µm. Fluorescent primary antibodies for microglial proliferation, synaptophysin, and postsynaptic density protein-95 (PSD95; also known as synapse-associated protein 90 (SAP90)) were purchased from Abcam and are described in Table 1. Brain sections were incubated with primary antibodies overnight at 4°C. Fluorescent images were captured using the BZ-X series KEYENCE All-in-One Fluorescence Microscope (Keyence, Itasca, IL, USA). Fluorescence signal intensity was analyzed using NIS-Elements BR3.10 software.

| Primary Antibodies |  |  |  |  |
| --- | --- | --- | --- | --- |
| Antigen | Host Species | Type | Conjugates | Catalogue Number |
| Iba1 | Rabbit | Monoclonal | Alexa Fluor 594 | ab178847 |
| Ki67 | Mouse | Monoclonal | Alexa Fluor 647 | ab279653 |
| Synaptophysin | Rabbit | Monoclonal | Alexa Fluor 488 | ab32127 |
| PSD95 | Rabbit | Polyclonal | Alexa Fluor 647 | ab18258 |

### Statistical Analysis

Statistical analyses were conducted using analysis of variance (ANOVA) and regression techniques (SPSS Statistics 28, IBM Corp., Somer, NY, USA; GraphPad Prism 5.02, GraphPad Software, Inc., La Jolla, CA). Figures were created using GraphPad Prism 5. An alpha level of *p*≤0.05 was established for statistical significance.

Univariate ANOVAs were conducted to evaluate genotype (EcoHIV vs. Control) induced alterations in microglia proliferation and synaptic dysfunction. The dependent variables investigated include the number of 1ba1+ cells expressing Ki67+ signals, an index of microglial proliferation, as well as the intensity of synaptophysin and PSD95, indices of synaptic dysfunction.

Multiple and simple regression analyses were utilized to evaluate the association between microglial proliferation and synaptic dysfunction. The dependent variable of interest for all regression analyses was the number of co-localized 1ba1+ and Ki67+ cells. First, a stepwise multiple regression approach was implemented, whereby the independent variables included previously published measures of neuronal and dendritic spine morphology in pyramidal neurons in the mPFC (Li et al., 2021a). More specifically, the number of dendritic branches at Branch Orders 2 through 10, the number of intersections at radii between 40 and 150 μm, and the number of dendritic spines within each head and neck diameter bin were included in the analysis. Second, the intensity of synaptophysin or PSD95 were used as dependent variables for two independent simple linear regressions.

## RESULTS

### Microglia are infected by HIV-1 viral proteins in the medial prefrontal cortex of HIV-1 seropositive individuals with HIV-1-associated neurocognitive disorders

Autopsy human brain tissues (*n*=9) of HIV-1 seropositive individuals with symptomatic HAND were provided by the National NeuroAIDS Tissue Consortium. A dual-labeling technique, whereby HIV-1 mRNA were visualized using RNAscope *in situ* hybridization (Figure 1A) and microglia were examined using immunostaining for Iba1 (Figure 1B), was implemented to investigate the cell-type expressing HIV-1 viral proteins in the dorsolateral prefrontal cortex (dlPFC; Brodmann’s Area 9) of HIV-1 seropositive individuals with synaptomatic HAND. Examination of the colocalization of HIV-1 mRNA and Iba1 (Figure 1C) supports microglia as the predominant cell-type to harbor HIV-1 viral proteins in the dlPFC; observations which confirm those previously reported by Ko et al. (2019). Additionally, given the potential role of activated microglia in neuronal dysfunction, synaptophysin, an indicator of presynaptic vesicle proteins, was examined (Figure 1D). Low levels of synaptophysin were observed in the dlPFC of HIV-1 seropositive individuals with symptomatic HAND.

**Figure 1.**
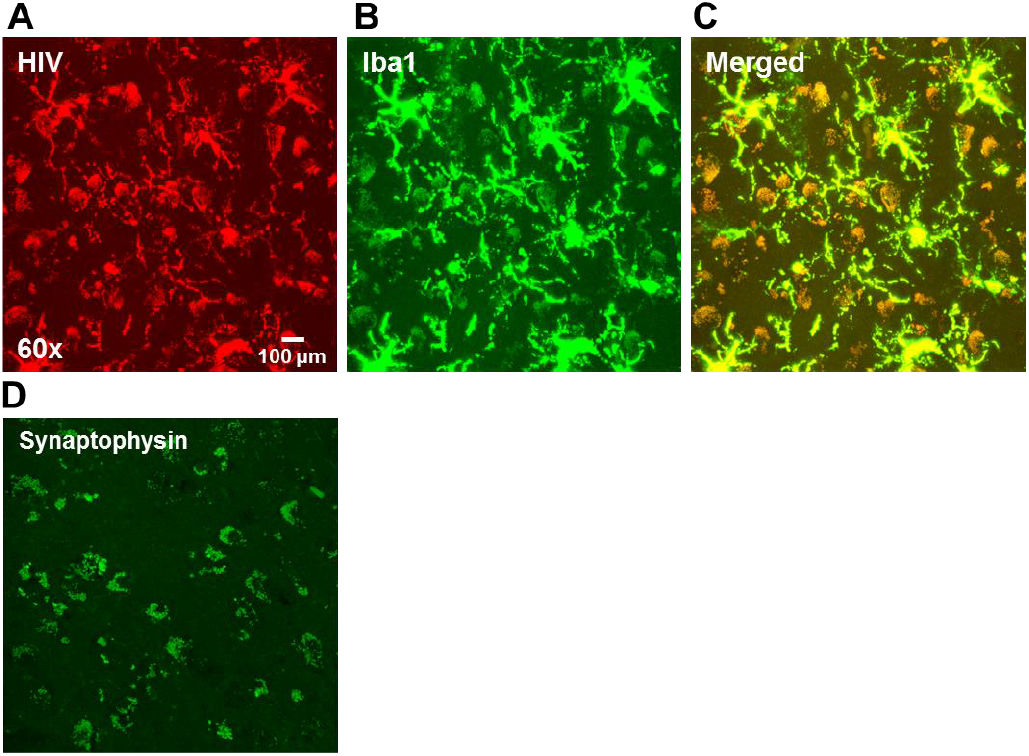
HIV-1 mRNA, microglial activation, and synaptophysin in the dorsolateral PFC (dlPFC) of HIV-1 seropositive individuals with symptomatic HIV-1-associated neurocognitive disorders (HAND). (**A-C**) Representative images illustrate the presence of pronounced HIV-1 mRNA (**A**), activation of microglia (**B**), and their co-localization (**C**) in the dlPFC. Observations support microglia as the predominant cell-type to harbor HIV-1 viral proteins in the dlPFC of HIV-1 seropositive individuals with symptomatic HAND. (**D**) A representative image demonstrates the overall low levels of synaptophysin present in HIV-1 seropositive individuals with HAND.

### Consistent with findings in HIV-1 seropositive individuals, EcoHIV infection in rats is harbored in microglia, which induces enhanced proliferation

A novel biological system to model systemic HIV-1 infection in the rat using chimeric HIV (EcoHIV) was recently reported (Li et al., 2021a; Li et al., 2021c). Consistent with findings in HIV-1 seropositive individuals with HAND, the EcoHIV rat harbors significant infection in microglia in the medial prefrontal cortex (mPFC) seven days (Li et al., 2021a) and eight weeks (Li et al., 2021a; McLaurin et al., 2022; Present Study, Figure 2A, 2B) post infection. In addition, EcoHIV-infected microglia in the mPFC appear activated, evidenced by an increased number of processes and larger soma under high magnification (Figure 2B).

**Figure 2.**
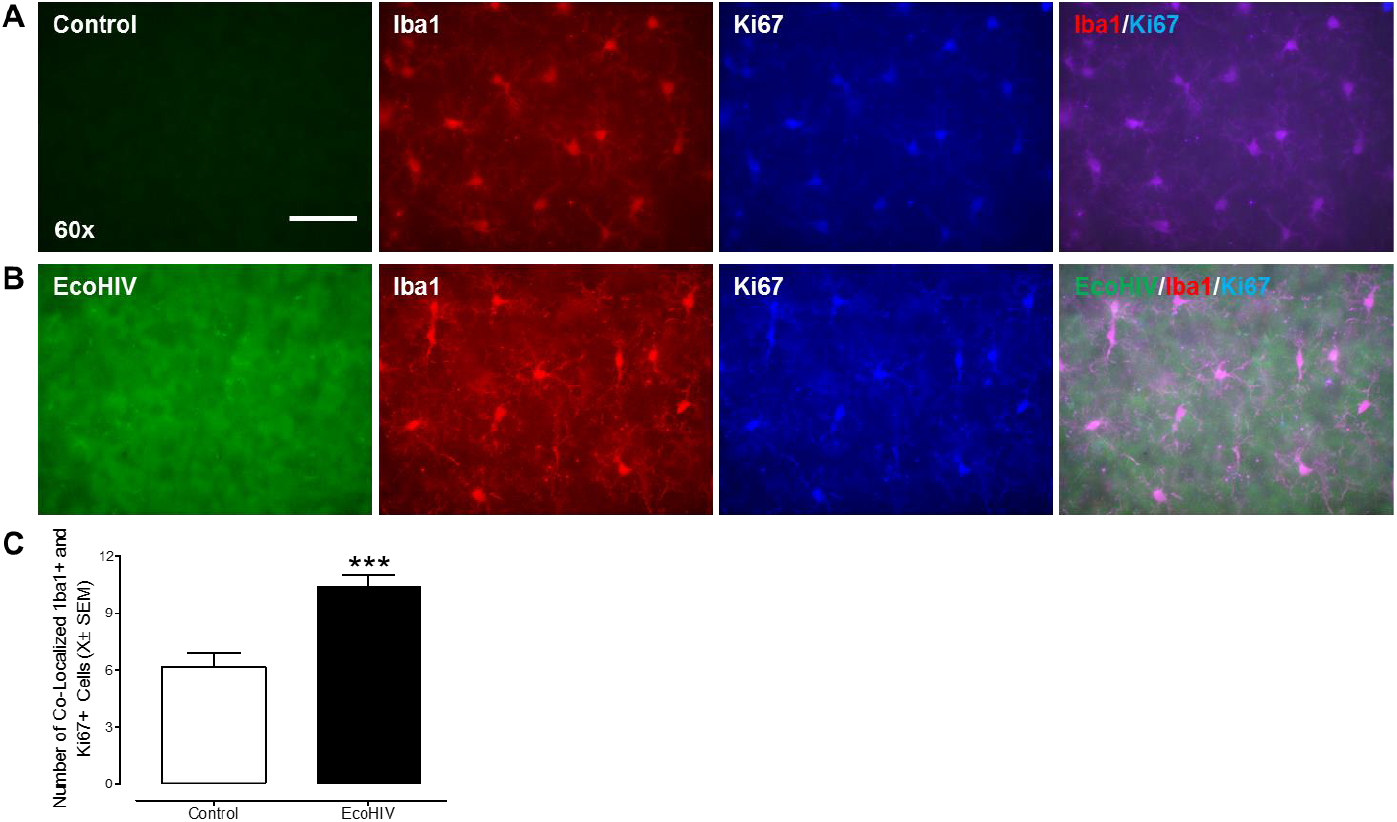
EcoHIV infection productively infected microglia, resulting in microglial activation and elevated proliferation. Brain tissue from the medial prefrontal cortex of control (**A**) and EcoHIV (**B**) rats were dual-labeled with a cell marker for microglia (i.e., Iba1) and proliferation (i.e., Ki67); EcoHIV EGFP was integral to the virus construct. Quantification of the number of co-localized Iba1+ and Ki67+ cells (**C**) revealed a statistically significant increase in EcoHIV, relative to control, animals.

To further evaluate and quantify how the active infection of microglia alters their proliferation, a dual-immunohistochemical approach was implemented, whereby brain tissue from the mPFC of control (*n*=6) and EcoHIV (*n*=6) animals was stained with Iba1^+^ and Ki67^+^ (Figure 2A,B). Quantification of the number of 1ba1+ cells expressing Ki67+ signals revealed a statistically significant increase in these cells in EcoHIV, relative to control, animals (Figure 2C; *F*(1,11)=20,0, *p*≤0.01). Taken together, EcoHIV productively infects microglia in the mPFC, resulting in their activation and elevated proliferation.

### Eight weeks after inoculation, EcoHIV animals exhibit pronounced synaptic dysfunction, evidenced by alterations in presynaptic and postsynaptic proteins

Profound neuronal damage (Moore et al., 2006) and synaptic dysfunction/loss (Gelman & Nguyen, 2010; Desplats et al., 2013; Weiss et al., 2021) have been purported as neural mechanisms underlying HAND in the post-cART era. Indeed, EcoHIV (*n*=6) animals, relative to control (*n*=6) animals, exhibit pronounced alterations in dendritic spines and neuronal morphology in pyramidal neurons in the mPFC (Li et al., 2021a). To examine the generalizability of these alterations in the EcoHIV rat, alterations in presynaptic vesicle proteins and postsynaptic scaffolding proteins were examined using synaptophysin and PSD95 markers, respectively. Quantification of the intensity of synaptophysin (Figure 3A-C) and PSD95 (Figure 3D-F) staining revealed statistically significant decreases in EcoHIV, relative to control, animals (Synaptophysin: *F*(1,11)=6.5, *p*≤0.03, η_p_^2^=0.393; PSD95: *F*(1,11)=18.1, *p*≤0.001, η_p_^2^=0.645).

**Figure 3.**
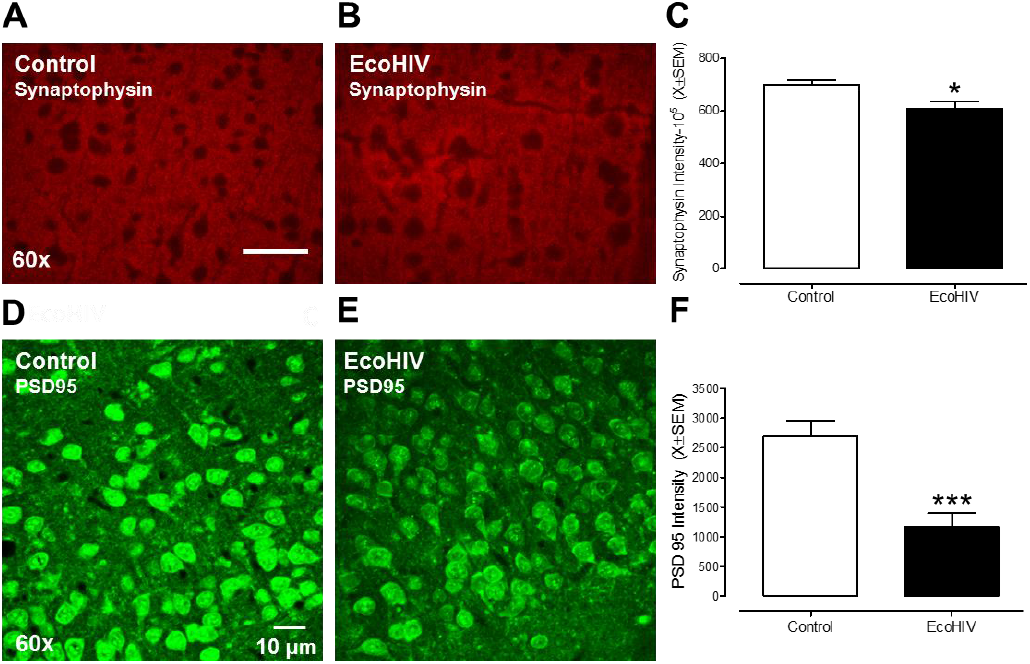
Animals inoculated with EcoHIV exhibited profound synaptic dysfunction in the medial prefrontal cortex. Representative images of synaptophysin (**A-B**), a presynaptic vesicle protein, and PSD95 (**D-E**), a postsynaptic scaffolding protein, are shown for both control and EcoHIV animals. Synaptophysin and PSD95 were quantified by examining the intensity of immunohistochemical staining, whereby both synaptophysin (**C**) and PSD95 (**F**) were significantly decreased in EcoHIV relative to control animals.

### Microglial proliferation may underlie synaptic dysfunction in EcoHIV rats

A multiple regression analysis was conducted to evaluate the relationship between microglia proliferation and previously published measures of neuronal and dendritic spine morphology (e.g., Sholl Analysis, Dendritic Spine Head and Neck Diameter) in pyramidal neurons in the mPFC (Li et al., 2021a). Indeed, 68.6% of the variance in the number of co-localized Iba1^+^ and Ki67^+^ cells was accounted for by two variables (Figure 4A,D; *F*(2,11)=9.8, *p* ≤ 0.005, *r*=0.828). Specifically, the two variables included in the regression model reflected the number of intersections at the 150 μm radius and the number of dendritic spines within the 0.60 μm neck diameter bin. Multiple regression analyses, therefore, support microglial proliferation as a potential mechanism underlying synaptic dysfunction.

**Figure 4.**
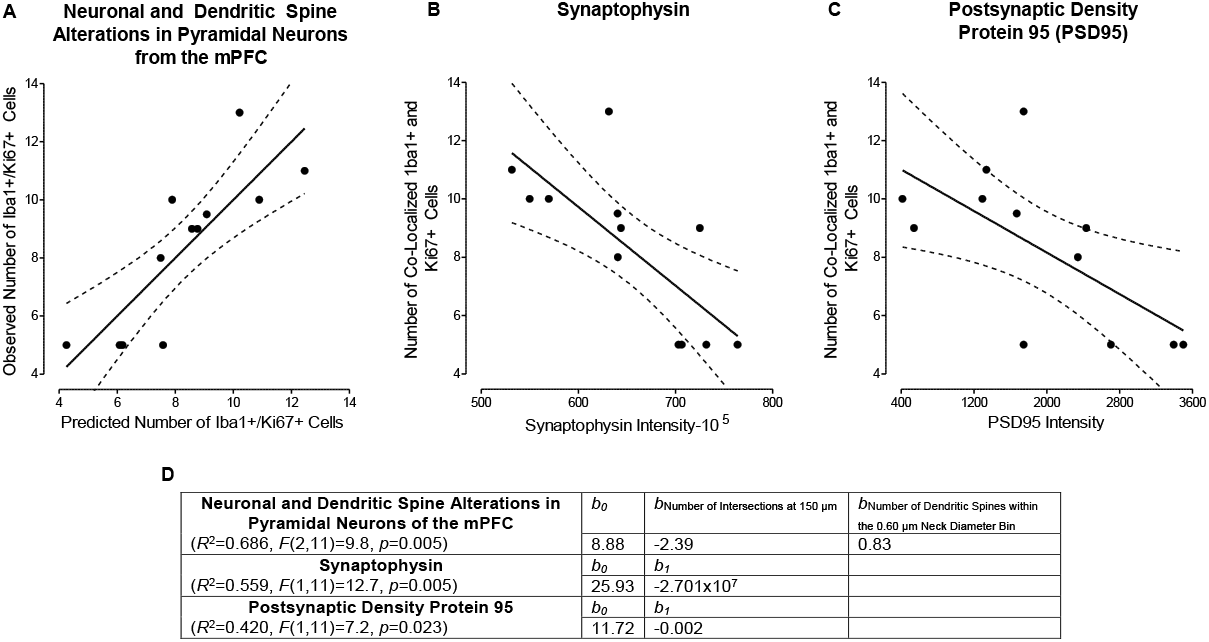
Regression analyses were conducted to evaluate the relationship between microglial proliferation and measures of synaptic dysfunction. (**A-C**) Microglial proliferation, indexed using the number of co-localized Iba1^+^ and Ki67^+^ cells, accounted for 68.6%, 55.9% and 42.0% of the variance in measure of neuronal and dendritic spine morphology, synaptophysin, and postsynaptic density protein 95, respectively. Statistical output from the regression analyses are shown in (**D**).

To further establish the generalizability of the relationship between microglial proliferation and synaptic dysfunction, simple linear regression analyses were conducted, whereby synaptic dysfunction was indexed using synaptophysin and PSD95. Microglial proliferation accounted for 55.9% of the variance in synaptophysin intensity (Figure 4B,D). Indeed, an increased number of co-localized Iba1^+^ and Ki67^+^ cells was significantly associated with lower synaptophysin intensity (*F*(1,11)=12.7, *p* ≤ 0.005, *r*=0.748). Further, 42.0% of the variance in the number of co-localized Iba1^+^ and Ki67^+^ cells was accounted for by intensity of PSD95 staining (Figure 4C-D; *F*(1,11)=7.2, *p* ≤ 0.023, *r*=0.648). Collectively, microglial proliferation accounts for a significant proportion of the variance in synaptic dysfunction affording a potential neural mechanism underlying HIV-1-associated neuronal and synaptic damage.

## DISCUSSION

Microglial proliferation induced by chronic HIV-1 viral protein exposure underlies synaptic dysfunction in the mPFC. Consistent with previous literature (e.g., Ko et al., 2019), HIV-1 mRNA in postmortem chronically infected HIV-1 seropositive individuals with symptomatic HAND colocalizes with the microglial marker Iba1. Critically, the infection of microglia in the EcoHIV rat recapitulates observations in HIV-1 seropositive individuals, affording a biological system to investigate the pathophysiology of HIV-infected microglia. Indeed, chronic EcoHIV infection induces enhanced microglial proliferation, as evidenced by an increased number of colocalized Iba1 and Ki67 cells. Synaptic dysfunction in the EcoHIV rat is characterized by neuronal and dendritic spine alterations (Li et al., 2021a), as well as pronounced decreases in presynaptic vesicle proteins (i.e., synaptophysin) and postsynaptic scaffolding proteins (i.e., PSD95). Fundamentally, regression analyses revealed the strong association (i.e., *r*=0.648-0.828) between microglial proliferation and three measures of synaptic dysfunction; an association which supports a neural mechanism underlying HIV-1-associated synaptodendritic alterations. Expanding upon our understanding of the pathophysiology of HIV-infected microglia and their role in the pathogenesis of HIV-1-associated neurocognitive and affective alterations affords key novel therapeutic targets.

Macrophage colony-stimulating factor (M-CSF; alternative acronym: CSF1) may underlie the profound microglial proliferation induced by chronic EcoHIV inoculation. Originally purified and characterized by Stanley and Heard (1977), M-CSF is a glycoprotein composed of two nearly identical subunits and a total molecular weight of approximately 70,000. M-CSF has been characterized as a strong microglial mitogen (Giulian & Ingeman, 1988) and is essential for posttraumatic microglial proliferation (Raivich et al., 1994; Berezovskaya et al., 1995). Indeed, the proliferation of microglia is induced by the actions of M-CSF on colony-stimulating factor-1 receptor (CSF-1R; also known as c-FMS; Gómez-Nicola et al., 2013). Fundamentally, M-CSF and/or its ligand CSF-1R is upregulated in multiple neurodegenerative disorders (e.g., Alzheimer’s disease: Akiyama et al., 1994, Walker et al., 2017; Prion Disease: Gómez-Nicola et al., 2013), including HIV-1 (Knight et al., 2018; Irons et al., 2019).

With regards to HIV-1, M-CSF plays a fundamental role in the pathogenesis of HIV-1 infection in both the periphery and CNS. Indeed, *in vitro* studies of macrophages cultured from peripheral blood have demonstrated that M-CSF signaling enhances the susceptibility of peripheral macrophages to infection (Kalter et al., 1991; Bergamini et al., 1994) and promotes viral replication (Kalter et al., 1991). In the CNS, chronically infected HIV-1 seropositive individuals on suppressive cART exhibit increased expression of CSF-1R relative to HIV-1 seronegative individuals. Consistent with clinical observations, elevated M-CSF mRNA and CSF-1R have been observed in the basal ganglia and/or frontal cortex of SIV-infected macaques; findings which develop early in the course of infection and persist despite cART (Knight et al., 2018). The fundamental importance of M-CSF upregulation is evidenced by the strong association between M-CSF and neurocognition, whereby higher CSF levels of M-CSF are associated with increased neurocognitive impairments in HIV-1 seropositive individuals (Lentz et al., 2010).

The profound, generalizable synaptic dysfunction observed in animals inoculated with EcoHIV may result, at least in part, from HIV-associated microglial proliferation. Indeed, synaptic dysfunction in EcoHIV rats is characterized by decreased expression of presynaptic vesicle proteins (i.e., synaptophysin; Present Study) and postsynaptic scaffolding proteins (i.e., PSD-95; Present Study), as well as morphological alterations in pyramidal neurons and associated dendritic spines from the mPFC (Previously Published, Li et al., 2021a). Synaptophysin, a glycoprotein whose molecular mass is 38,000 daltons, is an integral membrane protein that is enriched in the presynaptic vesicle (Jahn et al., 1985; Wiedenmann & Franke, 1985). Given that presynaptic vesicles are fundamental for the storage and release of neurotransmitters (for review, Murthy & De Camilli, 2003), decreased levels of synaptophysin in the EcoHIV rat supports a structural change underlying HIV-induced alterations in neurotransmission (e.g., Kumar et al., 2009; Denton et al., 2019). Further, PSD-95, a postsynaptic scaffolding protein that is member of the membrane-associated guanylate kinase (MAGUK) family, is essential for maintaining the molecular organization of the postsynaptic density (PSD; Chen et al., 2011). EcoHIV induced decreases in PSD-95, therefore, may afford a structural locus underlying HIV-1-associated changes to dendritic spine morphology (e.g., Roscoe et al., 2014; McLaurin et al., 2019; Li et al., 2021a). Indeed, a profound population shift towards a more immature dendritic spine phenotype (i.e., stubby; increased head and neck diameter) has been previously reported in EcoHIV rats (Li et al., 2021a; McLaurin et al., 2022).

Fundamentally, microglia are intricately involved in the development and maintenance of synaptic processes. Under homeostatic conditions, microglia are highly dynamic patrollers of the environment (Nimmerjahn et al., 2005), whereby their environmental surveillance is uniquely targeted to synaptic structures (Wake et al., 2009; Tremblay et al., 2010). Indeed, during early development, microglia modulate synapse formation (Miyamoto et al., 2016) and selectively eliminate excess axons and synapses (e.g., Paolicelli et al., 2011; Weinhard et al., 2018; Lim & Ruthazer, 2021) via direct contact. The fundamental role of microglia persists into adulthood, whereby microglia are involved in synapse phagocytosis during sleep (Choudhury et al., 2020) and aging (Tremblay et al., 2012), cellular communication (Antonucci et al., 2012; Akiyoshi et al., 2018), and cognitive processes (Torres et al., 2016).

Given the intricate relationship between microglia and synaptic processes, simple linear and multiple regression analyses were utilized to establish the relationship between microglia proliferation and synaptic dysfunction. Microglial proliferation was significantly associated with measures of neuronal and dendritic spine morphology, synaptophysin, and PSD-95, whereby it explained 68.6%, 55.9% and 42% of the variance in these synaptic indicators, respectively. It is notable, albeit unsurprising, that microglial proliferation is most strongly (*r*=0.828) related to measures of neuronal and dendritic spine morphology; a finding which may be due to two factors. First, multiple regression analyses were conducted affording an opportunity to include several independent variables in the analysis. Measures reflecting dendritic branching complexity (i.e., Branch Order), dendritic arbor complexity (i.e., Sholl Analysis), and dendritic spine morphology (i.e., Head Diameter, Neck Diameter) were included in the analysis. Second, three-dimensional reconstruction of pyramidal neurons, and associated dendritic spines, provides a wealth of data that reflects multiple aspects of neuronal functioning. For example, the population shift towards a stubby dendritic phenotype in EcoHIV rats supports decreased synaptic strength, as recent data demonstrates that the stubby, relative to mushroom, spine proteome exhibits a lower association with synaptic strength (Helm et al., 2021). In addition, dopamine receptors are concentrated on the dendritic spine neck, whereas glutamatergic afferents establish contact primarily with receptors on the dendritic spine head (Goldman-Rakic et al., 1989; Carr & Sesack, 1996); the absence of a dendritic spine neck (as in stubby spines), therefore, suggests decreased dopaminergic innervation. Investigation of neuronal and dendritic spine morphology via three-dimensional reconstruction analysis, therefore, provides a more comprehensive evaluation of how constitutive expression of HIV-1 viral proteins alters both the structure and function of neurons.

Taken together, the present study enhances our understanding of the relationship between microglial and synaptic dysfunction resulting from chronic HIV-1 viral protein exposure. EcoHIV rats harbor HIV-1 mRNA in microglia resembling HIV-1 seropositive individuals with HAND affording a novel biological system to investigate HIV-1-associated synaptic dysfunction. Indeed, EcoHIV rats exhibited profound microglial proliferation, as well as generalizable synaptic dysfunction. Fundamentally, microglial proliferation is significantly associated with three measures of synaptic dysfunction. Expanding upon our understanding of the pathophysiology of HIV-infected microglia and their role in the pathogenesis of HIV-1-associated neurocognitive and affective alterations affords key novel therapeutic targets.

## CONFLICT OF INTEREST

The authors declare that they have no conflict of interest.

## ACKNOWLEDGEMENTS

This work was supported in part by grants from NIH (National Institute on Drug Abuse, R01-DA013137; National Institute on Drug Abuse, K99-DA056288, National Institute of Mental Health, R01-MH106392; National Institute of Neurological Disorders and Stroke, R01-NS100624).

## Notes

### Competing Interest Statement

The authors have declared no competing interest.

